# Zika in Rio de Janeiro: Assessment of basic reproduction number and comparison with dengue outbreaks

**DOI:** 10.1101/055475

**Authors:** D. A. M. Villela, L. S. Bastos, L. M. de Carvalho, O. G. cruz, M. F. C. Gomes, B. Durovni, M. C. Lemos, V. Saraceni, F. C. Coelho, C. T. Codeço

## Abstract

Zika virus infection was declared a public health emergency of international concern in February 2016 in response to the outbreak in Brazil and its suspected link with congenital anomalies. In this study we use notification data and disease natural history parameters to estimate the basic reproduction number (*R*_0_) of Zika in Rio de Janeiro, Brazil. We also obtain estimates of *R*_0_ of dengue from time series of dengue cases in the outbreaks registered in 2002 and 2012 in the city, when DENV-3 and DENV-4 serotypes respectively, had just emerged. Our estimates of the basic reproduction number for Zika in Rio de Janeiro based on surveillance notifications (*R*_0_ = 2.33, 95% CI: 1.97 − 2.97) were higher than those obtained for dengue in the city (year 2002: *R*_0_ = 1.70 [1.50 − 2.02]; year 2012: Ro = 1.25 [1.18 − 1.36]). Given the role of *Aedes aegypti* as vector of both the Zika and dengue viruses, we also derive Ro of Zika as a function of both dengue reproduction number and entomological and epidemiological parameters for dengue and Zika. Using the dengue outbreaks from previous years allowed us to estimate the potential *R*_0_ of Zika. Our estimates were closely in agreement with our first Zika’s *R*_0_ estimation from notification data. Hence, these results validate deriving the potential risk of Zika transmission in areas with recurring dengue outbreaks. Whether transmission routes other than vector-based can sustain a Zika epidemic still deserves attention, but our results suggest that the Zika outbreak in Rio de Janeiro emerged due to population susceptibility and ubiquitous presence of *Ae. aegypti.*

## Introduction

In February 2016, Zika virus infection was declared a public health emergency of international concern [1] in response to the outbreak in Brazil and its suspected link with congenital anomalies [2, 3, 4]. This came as a surprise, as since its first isolation in 1947 in the Zika forest in Uganda [5], the virus was associated with benign disease and had remained mostly unnoticed. Isolated outbreaks have been reported before in Africa and Asia/Oceania but all of them involved small populations [6]. In Brazil Zika virus (ZIKV) found a large pool of susceptible individuals, and the range of possible clinical outcomes became apparent, including birth defects, neurological and auto-immune disorders [3]. According to outbreak reports [7], the epicentre of the American epidemic was in North-East Brazil, where ZIKV was introduced in mid 2014, although a molecular study [8] suggested introduction took place in 2013. In 2016, all 26 Brazilian states had confirmed local transmission of ZIKV. In the Americas, 34 countries and territories had already confirmed autochthonous ZIKV cases by April 2016.

Transmission of ZIKV to humans is mostly attributed to mosquitoes of the genus *Aedes (Stegomyia*). Infected mosquitoes were found in localities reporting outbreaks, in Gabon *Aedes albopictus* mosquitoes and in Yap Island *Aedes hensilii* mosquitoes [6]. *Ae. aegypti* mosquitoes are widely suspected to be the primary vector in urban centres in the Americas, based on its widespread distribution and role as dengue vector. In the recent Zika epidemic in Rio de Janeiro, *Aedes aegypti* mosquitoes were found naturally infected with Zika virus [9].

Rio de Janeiro is among the most dengue affected cities in Brazil. Rio is a large urban centre with 6.5 million inhabitants which was the port-of-entry in the country of three of four current circulating dengue viruses. Climatic and environmental conditions favour year-round transmission of dengue, with a well characterized seasonal profile. The city has also recently been in the international spotlight due to major sporting events, including 2014 FIFA World Cup and the 2016 Summer Olympic and Paralympic Games.

Quantitative knowledge about the transmission risk of arboviruses is vital to disease surveillance. Estimation of the basic reproduction number, *R*_0_, provides a measurement of the transmission potential of the virus. Such measurement is important to support preparedness plans and to assess risk of epidemic emergence into disease-free areas. Moreover, estimation of *R*_0_ can also contribute to the understanding of the epidemiology of this disease and how it changes geographically and temporally.

Here, we report estimates of *R*_0_ of Zika in Rio. We apply two methods for estimation of Zika’s basic reproduction number, which mainly differ by whether or not they use Zika notification data. First, we estimate *R*_0_ from Zika notification cases in the city of Rio de Janeiro in 2016. For comparison with dengue epidemics in the same area in previous years, we also estimate the basic reproduction number of dengue using notification cases from two outbreaks occurring just after introductions of dengue serotypes 3 and 4 in the city [10, 11]. We then derive the expected basic reproduction number of Zika as function of the *R*_0_ estimates for dengue and compare these estimates with the ones obtained from notification data. Our results show that estimating *R*_0_ for Zika using information for dengue yields estimates in agreement with estimates obtained directly from Zika notification data. This validates using information on dengue to gain insight into the risk of Zika transmission prior to the introduction of the virus in an at-risk area with presence of the vector.

## Methods

### Data

Infectious disease surveillance in Brazil is handled by the Brazilian Notifiable Diseases Information System (SINAN), where each suspected, and eventually confirmed, case of Zika infection is notified as ICD-10 diagnosis code A92.8 (other specified mosquito-borne viral fevers). Dengue surveillance in Brazil dates back to the 1990s, and suspected cases are notified in the SINAN as ICD-10 diagnosis code A90 (Dengue fever) or A91 (Dengue hemorrhagic fever). Dengue serotype DENV-3 was first observed in the state of Rio de Janeiro, in the neighbouring Nova Iguaçu city, in January 2001 [10], while the serotype DENV-4 was first observed in Rio de Janeiro state in Niterói city in March 2011 [11].

In order to calculate dengue’s reproduction number, we assume the epidemics of 2002 and 2012 in the city of Rio de Janeiro were mainly caused by serotypes DENV-3 and DENV-4, respectively. These are taken to represent the introduction of new serotypes, for which the population had no previous immunity. This is important in order to make estimates comparable with Zika.

Case notification time series of dengue and Zika were constructed by aggregation by epidemiological week. Both Zika and dengue notification data are stratified by ten health districts (HD), which are essentially health surveillance sub-areas in the city of Rio de Janeiro.

#### Estimation of exponential growth rate from the epidemic curve

In order to estimate the weekly exponential rate of the epidemic curve, given by the number of cases infected by Zika, we fitted a linear model of the logarithm of notification counts adjusted by time, given by the number of weeks of the early outbreak period. We take the 43rd epidemic week of 2015 (from the 18th to the 24th of October 2015) as the starting week, after which notification of suspected cases of Zika became mandatory in the city of Rio de Janeiro. In order to select the end of the early outbreak period we apply the time windows which minimise the sum of residuals.

We also estimated the exponential rates for each one of the ten health districts (HD) of Rio de Janeiro, using a mixed linear model, where the number of reported cases in each HD is proportional to 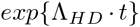, with Λ_*HD*_ = Λ_0_ + *λ*_HD_. Parameter Λ_0_ is a baseline rate, and *λ*_*HD*_ is a zero-mean random effect by district. We use the same early outbreak period defined in the overall case in order to estimate the exponential rate by health district.

#### Estimation of the basic reproduction number (*R*_0_) from notification data

We apply the *R*_0_ formulation proposed by Pinho *et al.* [12] to model the dynamics of dengue fever to assess both Zika’s and dengue’s basic reproduction number in the city of Rio de Janeiro from SINAN data collected from described outbreaks of these diseases. The model considers vector-borne transmission, by defining compartments of susceptible, exposed and infected mosquitoes, and for humans, susceptible, exposed, infected and recovered. Hence, the disease transmission involves a cycle of two infectious generations, mosquitoes and humans. The concept of basic reproduction number gives us the average number of secondary cases per generation after an initial infected individual.

This approach relies on the assumption that the number of cases in the early outbreak grows exponentially, hence proportional to exp{Λ·*t*}, where *t* is the time in weeks since the outbreak start and Λ is the exponential growth rate of the number reported cases. An estimate of the basic reproduction number is given by the following equation:

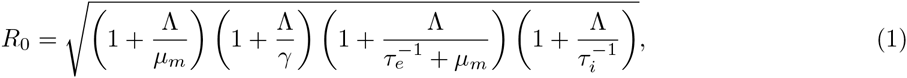

where *γ* is the human recovery rate, *μ*_*m*_ is the mosquito mortality rate, *τ*_*i*_ is the median intrinsic incubation period in humans, and *τ*_*e*_ is the median extrinsic incubation period in mosquitoes. In equation 1 we neglect adult mosquito control, *c*_*m*_, as well as human mortality rate, *μ*_*h*_, which are present in the original formula in [12]. The former is taken to be zero since no structured intervention was taking place during the time window analysed. Regarding the latter, human mortality rate in Brazil is orders of magnitude lower than the intrinsic incubation period and human recovery rate, which are on the order of a few days. The life expectancy at birth in Brazil was of 75 years in 2014. Therefore, we can safely neglect the human mortality rate *μ*_*h*_ for our purposes of estimating *R*_0_. We compiled a range of values for the necessary parameters taken from previous studies in Table 1.

**Table 1.**
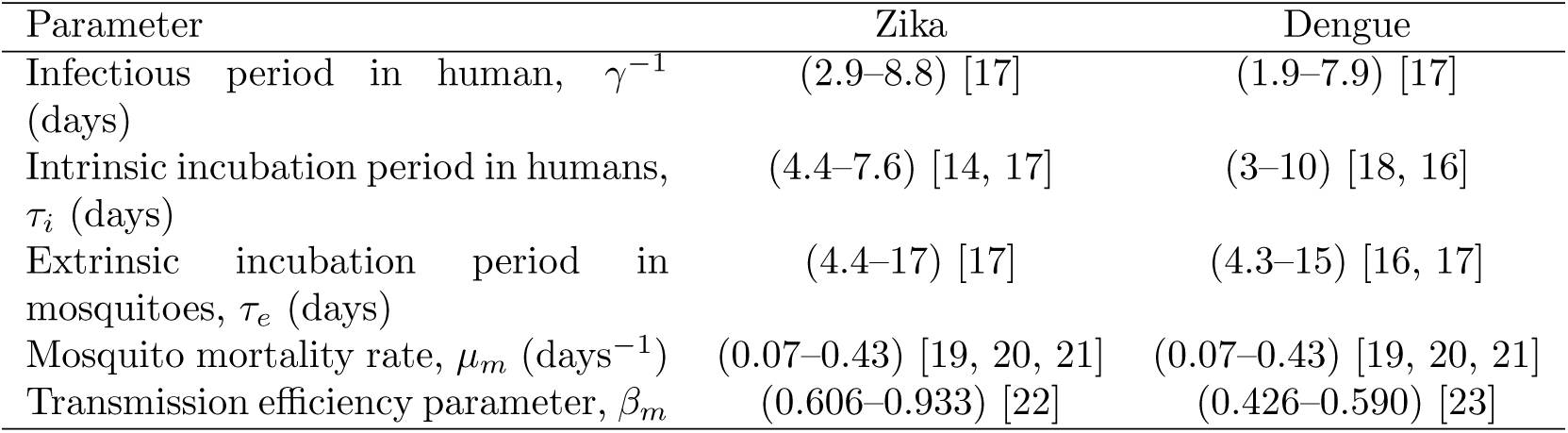
Zika and Dengue fever natural history parameters collated from the literature.

| Parameter | Zika | Dengue |
| --- | --- | --- |
| Infectious period in human, $\gamma^{-1}$ (days) | (2.9–8.8) [17] | (1.9–7.9) [17] |
| Intrinsic incubation period in humans, $\tau_i$ (days) | (4.4–7.6) [14, 17] | (3–10) [18, 16] |
| Extrinsic incubation period in mosquitoes, $\tau_e$ (days) | (4.4–17) [17] | (4.3–15) [16, 17] |
| Mosquito mortality rate, $\mu_m$ (days$^{-1}$) | (0.07–0.43) [19, 20, 21] | (0.07–0.43) [19, 20, 21] |
| Transmission efficiency parameter, $\beta_m$ | (0.606–0.933) [22] | (0.426–0.590) [23] |

#### Potential basic reproduction number of Zika

Massad *et al.* [13] derived a mathematical method for estimating the reproduction number of yellow fever indirectly using an estimation of dengue’s basic reproduction number obtained using the exponential growth method. The underlying assumption was that both diseases share the same vector, and consequently, some parameters of their *R*_0_ expressions are the same. For instance, this approach does not require knowledge of the density of mosquitoes, which is usually hard to estimate.

We use this rationale to derive an expression for Zika’s basic reproduction number in a dengue endemic area, assuming that mosquitoes bite at the same rate and survive with the same daily probability, regardless of the virus they are infected with. From the *R*_0_ derivation found by Pinho *et al.* [12] for their model, we have the following expressions for the reproduction number of Zika and dengue, *R*_0, *z*_ and *R*_0, *d*_, respectively:

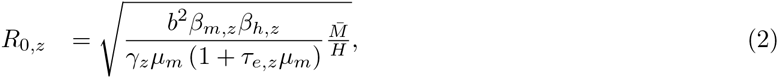

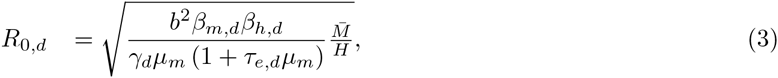

where the total mosquito population size is denoted by 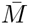, the human population size by *H*, the mosquito biting rate *b*, the proportion *β*_*m*_ of mosquito bites in infected humans considered to be infective to the vector, and the proportion *β*_*h*_ of infected mosquito bites effectively infective to humans - parameters are indexed by dengue (*d*) and Zika (*z*), accordingly. Some of these parameters are difficult to estimate, but by taking the ratio between equations 2 and 3, we obtain R_0*z*_ indirectly from R_0,*d*_:

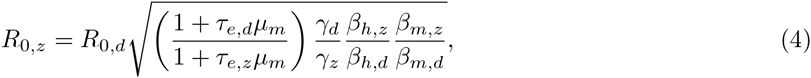

which is convenient because some parameters are cancelled out. If one assumes that bites from mosquitoes infected with any of these two viruses are equally likely to infect susceptible humans, that is *β*_*h,z*_ = *β*_*h,d*_, it is possible to estimate the basic reproduction rate of Zika based on that of dengue obtained from previous epidemics, assuming infection routes are the same. On the other hand, by estimating *R*_*z*_ and *R*_*d*_ independently (direct approach), one can use equation 4 to estimate the ratio *β*_*h,z*_/*β*_*h,d*_ if the remaining parameters are known, still assuming that transmission dynamics are identical.

Hence, this method requires a time series of dengue cases and knowledge of entomological parameters describing the vectorial competence, incubation period, and human recovery rate. Such estimation can be applied to areas that have experienced dengue outbreaks with potential to develop a Zika epidemic.

From the two methods used in this work, the one proposed by Pinho *et al.* [12] assumes the number of cases to exhibit an exponential increase, hence *R*_0_ > 1. However, the second method might theoretically yield *R*_0_,_*z*_ < 1, depending on infectivity parameters, incubation periods and recovery rates of both dengue and Zika even if the number of dengue cases is assumed to grow exponentially (i.e. *R*_0,*d*_ > 1).

#### Parameter uncertainty

The use of the equations (1) and (4) requires knowledge about the disease natural history parameters. Table 1 presents a compilation of the literature on necessary parameters to calculate *R*_0_ according to different methods. A systematic review of the literature on Zika [14] published estimates of incubation and infection periods of ZIKV based on (only) 25 Zika cases, mostly among Europeans and north Americans returning from Zika endemic countries and found values consistent with dengue [12, 15, 16]). Another study [17] comparing outbreaks of Zika in the Pacific islands of Micronesia, the Yap Main Islands and Fais, has found similar incubation and infection periods for Zika. The mosquito mortality rate is obtained from various reports [19, 20, 21] on mark-release-recapture experiments with *Ae. aegypti* mosquito population in Rio de Janeiro which varies widely depending on different urban landscapes.

We assume that the uncertainty of each natural history parameter is represented by a Gaussian distribution whose mean and standard deviation are calculated based on the values presented in Table 1 where each interval is assumed to be a symmetric 99% probability interval. The uncertainty for exponential rate of case numbers is also represented by a Gaussian distribution, where the mean is given by the maximum likelihood estimate (MLE), 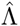, and the standard deviation is given by the observed standard deviation of 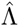. Hence, we can derive the induced distribution of *R*_0_ based on equations (1) and (4) using a Monte Carlo algorithm.

### Results

From January 2015 to mid-April 2016, 25, 213 suspected cases of Zika were notified for the city of Rio de Janeiro. Of these, 17, 585 were women and 7, 628 were men, what yields an attack ratio of approximately 395 per hundred thousand inhabitants over the entire period. Figure 1 shows the time series of weekly incidence in the city.

**Figure 1:**
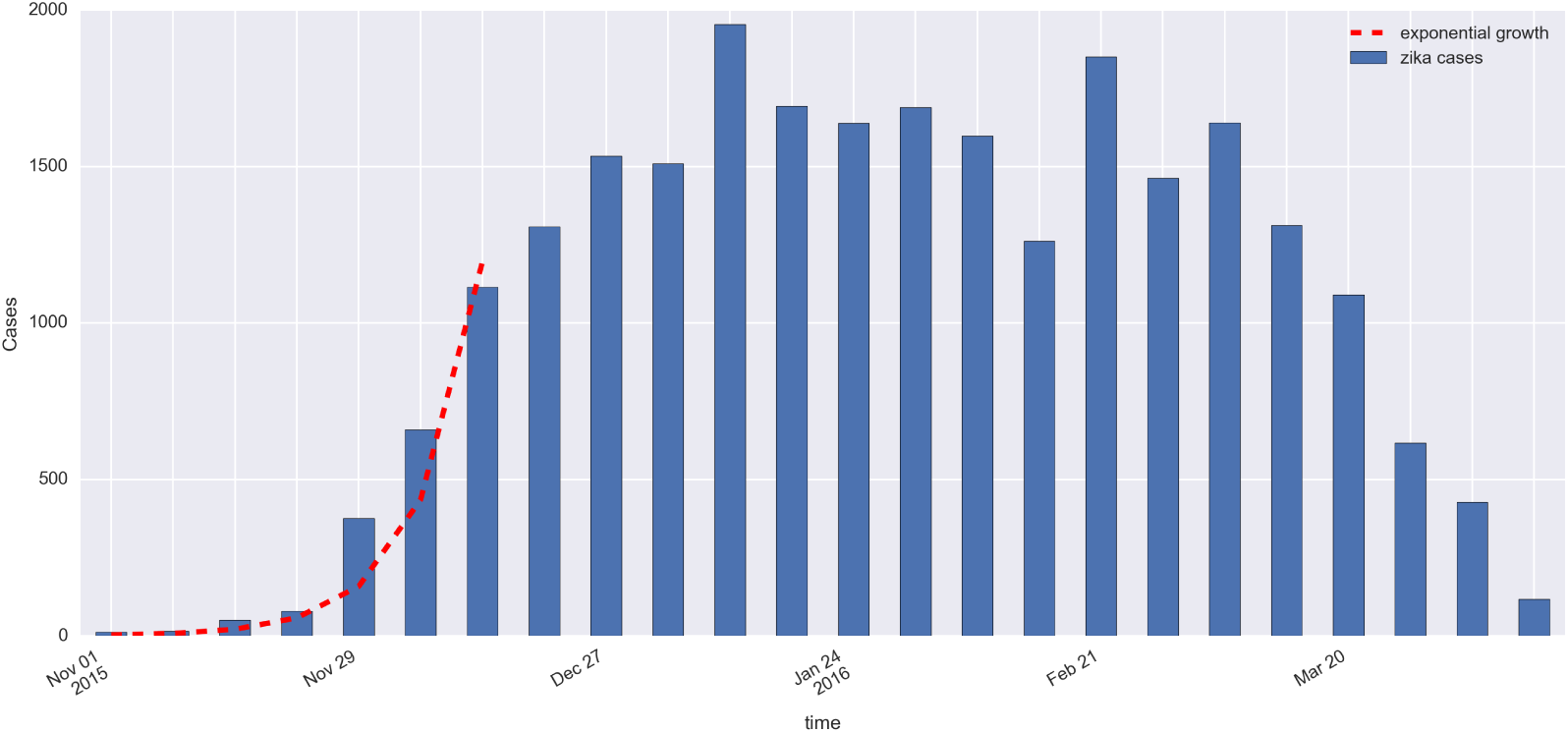
Epidemic curve of Zika in Rio de Janeiro, 2015-16 (blue bars). Red dashed line shows the exponential growth of Zika cases, with an estimated constant rate 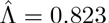, during the first seven weeks.

In Table 2, we vary the total number of weeks used to estimate Λ, and based on goodness-of-fit statistics for a linear model, the optimal value for the number of weeks is 7 weeks. The city-wide estimated rate Λ for Zika was 0.823 per week with standard deviation of 0.053.

**Table 2:** Estimates of exponential rate Λ for different numbers of weeks using the following linear model: *log*(*Y*_t_) = *β*_0_ + Λ_*t*_, where *Y* for the number of notified cases at week *t*; *t* = 1, 2,…, *T*. The first week, *t* = 1, is the 43th epidemic week of 2015 (from the 18th to the 24th of October 2015), *β*_0_ is the intercept and the coefficient Λ is the force of infection. The residual standard deviation, σ and the coefficients of determination, *R*^2^ and adjusted *R*^2^ were calculated each adjusted model.

| Total number of weeks ($T$) | Intercept | $\hat{\Lambda}$ | $\sigma$ | $R^2$ | adjusted $R^2$ |
| --- | --- | --- | --- | --- | --- |
| 6 | 1.305 | 0.852 | 0.296 | 0.973 | 0.966 |
| 7 | 1.381 | 0.823 | 0.279 | 0.980 | 0.976 |
| 8 | 1.583 | 0.756 | 0.358 | 0.969 | 0.964 |
| 9 | 1.820 | 0.685 | 0.460 | 0.950 | 0.943 |
| 10 | 2.091 | 0.611 | 0.579 | 0.920 | 0.910 |
| 11 | 2.312 | 0.556 | 0.640 | 0.902 | 0.891 |

We estimate the basic reproduction number for ZIKV in Rio de Janeiro from notication data at *R*_0_ = 2.33 (95% Confidence Interval: 1.97–2.97). Table 3 presents the estimates for the basic reproduction number for Zika by health district in Rio de Janeiro. The map of Rio de Janeiro (Figure 2) is also depicted in which health districts are shown by the estimated *R*_0_ for Zika. Health districts 3.2, 3.3 and 5.2 were those with highest reproduction numbers, all *R*_0_ being greater than four on average. Also, these areas are historically the areas with more notified cases of Dengue. Estimates of the basic reproduction number *R*_0_ for dengue from notification data in the years of entrance of DENV-3 and DENV-4 (2002 and 2012) in Rio de Janeiro are *R*_0_ = 1.70 (95% CI: 1.50-2.02) and *R*_0_ = 1.25 (95% CI: 1.18–1.36), respectively.

**Table 3:** Estimates for the basic reproduction number (*R*_0_) for Zika by health district in Rio de Janeiro.

| Health district | $\hat{R}_0$ | 2.5% | 97.5% |
| --- | --- | --- | --- |
| 1.0 | 2.09 | 1.78 | 2.54 |
| 2.1 | 1.91 | 1.54 | 2.27 |
| 2.2 | 1.98 | 1.68 | 2.31 |
| 3.1 | 2.10 | 1.91 | 2.35 |
| 3.2 | 2.35 | 2.01 | 2.80 |
| 3.3 | 2.44 | 2.15 | 2.74 |
| 4.0 | 2.12 | 1.90 | 2.54 |
| 5.1 | 2.02 | 1.74 | 2.47 |
| 5.2 | 2.55 | 2.18 | 3.07 |
| 5.3 | 2.15 | 1.94 | 2.42 |

**Figure 2:**
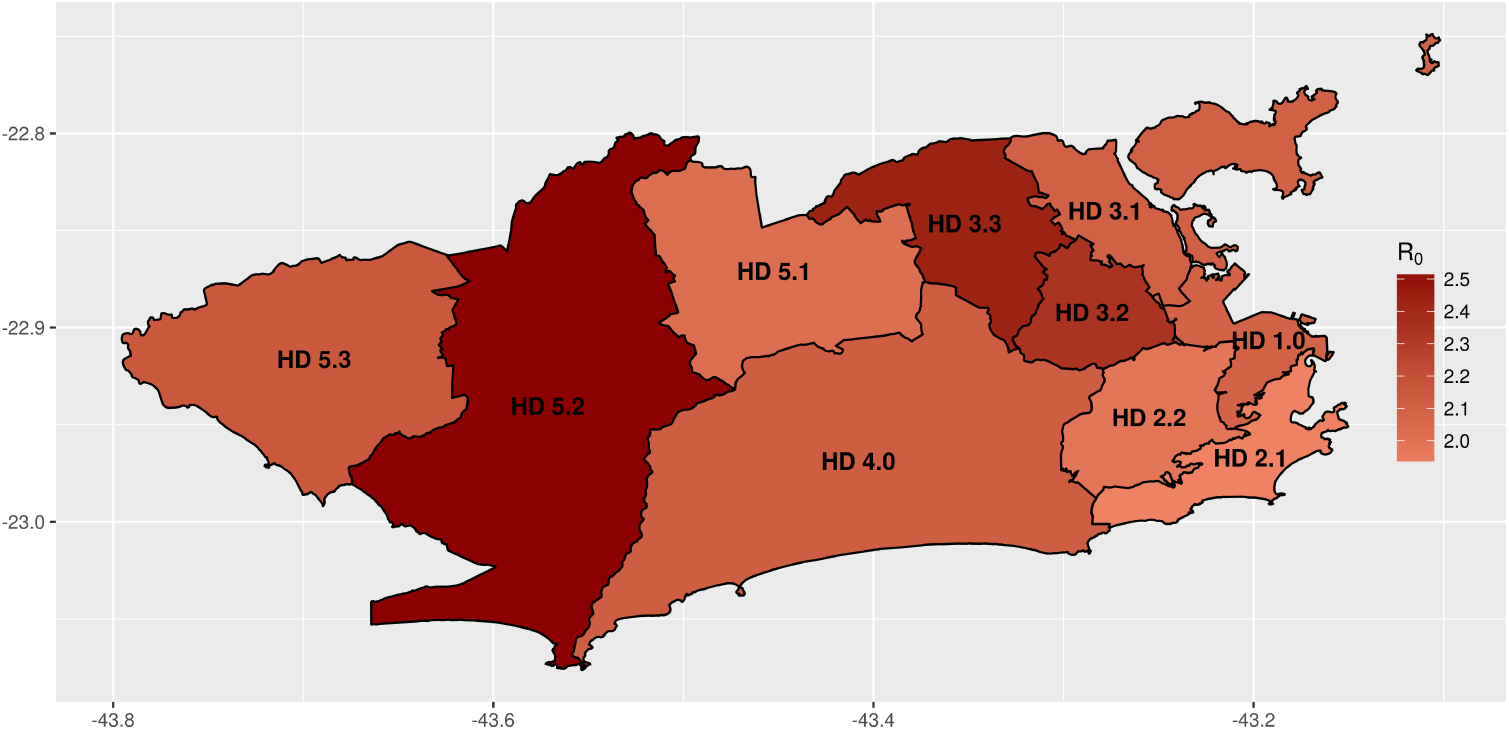
Estimates of the basic reproduction number for Zika by health districts in Rio de Janeiro.

Applying the disease natural history parameters from Table 1 into equation (4) indicates the *R*_0_ for Zika should be 1.4 times greater than the reproduction number for dengue. Hence, we estimate Zika’s basic reproduction number by the second method to find *R*_0_ = 2.45 (95% CI: 1.57-3.65) using our estimate of reproduction number of dengue in 2002, and *R*_0_ = 1.82 (95% CI: 1.19–2.68) using our estimated number from 2012. The estimates obtained using the first method are within these intervals. Table 4 summarises the basic reproduction numbers found for Zika in the 2016 epidemic, for dengue in 2002 and 2012 and the potential Zika *R*_0_ using such dengue numbers.

**Table 4:** Estimates and 95%CI for the basic reproduction number for Zika and dengue in Rio de Janeiro. We report estimates for Zika in 2015 and dengue in 2002 and 2012 obtained using equation (1). Estimates of Zika’s basic reproduction number using dengue data are obtained using equation (4) (see Methods)

| | $\hat{R}_0$ | 2.5% | 97.5% |
| --- | --- | --- | --- |
| Zika | 2.33 | 1.97 | 2.97 |
| Dengue 2002 | 1.70 | 1.50 | 2.02 |
| Dengue 2012 | 1.25 | 1.18 | 1.36 |
| Zika using dengue 2002 | 2.45 | 1.57 | 3.65 |
| Zika using dengue 2012 | 1.82 | 1.19 | 2.68 |

## Discussion

The basic reproduction number of Zika in Rio de Janeiro was estimated at 2.33 (95% CI: 1.97–2.97) given the number of notifications collected at the SINAN database. This value is consistent with estimates for French Polynesia, which varied from 1.5 to 3.1 [24, 25], but lower than those calculated for the Yap Island, ranging from 4.5 to 5.813 [24].

Estimates of Zika reproduction number in countries of Latin America reported by Ferguson *et al.* [26] have been used to obtain projection scenario and evaluate intervention policies. This study used a methodology different than ours, since they evaluate the reproduction number over time for several countries, including Brazilian states. We focus here on Zika and dengue notification data for the city of Rio de Janeiro to provide a comparison between reproduction numbers in local outbreaks and the basic reproduction number of Zika for multiple areas across the city. We also evaluate a method to calculate *R*_0_ without notification data, using *R*_0_ estimates for dengue and disease natural history parameters and find that this method yields estimates consistent with those obtained directly from notification data. The estimate of the reproduction number by Ferguson *et al.* [26] in the early 2016 epidemic in the state of Rio de Janeiro was in the range from 1.7 to 2.2, hence close to our estimates for the Zika outbreak in the city, after we adjusted the state estimates to be consistent with using *R*_0_ by generation, since a human-to-human reproduction number was reported (square root analysis).

Our estimates for Zika are also within the range of other viral diseases transmitted by *Aedes aegypti,* as dengue and Chikungunya, as reported in the literature [27]. Dengue serotypes 3 and 4 were introduced in Rio de Janeiro in 2002 and 2012, respectively. Assuming the population was mostly naive to these viruses allows us to compare Zika’s basic reproduction number to dengue’s *R*_0_ within the same city under an invasion scenario. Zika’s *R*_0_ in 2015 was 1.4 times greater than that of dengue epidemic of 2002, just after introduction of DENV-3, and 1.9 times greater than the estimation for dengue epidemic in 2012, just after introduction of DENV-4. These differences in transmission can be attributed to several factors. For example, since DENV-3 and DENV-4 invaded a population where the remaining dengue viruses were already circulating, some immunological protection could already be in place. It is also possible that vector competence differed between strains, either due to ecological, physiological, and genetic mechanisms [28] or year-specific (seasonal) factors [29].

The first wave of the Zika epidemic in Rio de Janeiro showed exponential growth during 7 weeks, and plateau at around 1500 cases per week during the whole summer of 2015-2016 (Figure 1). During this same period, dengue notification increased from 200 cases per week in November to 500 in February 2016 and to 1000 in April 2016 (http://info.dengue.mat.br). This growth in dengue cases, associated with high temperatures observed in this El Nino year, indicate that conditions for mosquito borne transmission existed. Forested mountains cross the city and create micro-climates in the valleys were most of the population lives; population density and living conditions also vary widely across the city. When calculating *R*_0_ at the health district level, we find that mean *R*_0_ varies from 1.91 to 2.55, indicating some level of heterogeneity. Understanding spatial variation is important, as those who live far apart are less likely to infect one another than those who live in closer proximity to each other. Spatial heterogeneity is known to slow down epidemics and this could be an explanation for the pattern observed [30].

A report by Chouin-Carneiro *et al.* [31] on the transmission efficiency of Zika virus using a ZIKV strain from New Caledonia suggested a low efficiency by *Ae. aegypti* mosquitoes. Considering the low transmission efficiency reported by Chouin-Carneiro *et al.* [31] a much lower Zika reproduction number would be expected, not excluding that *R*_0_ < 1. Conversely, Fernandes *et al.* [22] used combinations of Zika virus strains isolated in the city of Rio de Janeiro and a larger number of mosquitoes from local populations. This study found a much higher competence for *Ae. aegypti* to transmit ZIKV compared to results by Chouin-Carneiro *et al.* and also found no competence of *Culex* mosquitoes. Since a Zika epidemic was indeed observed in the city, a high competence by *Ae. aegypti* is a reasonable explanation, but more studies are recommended towards a more definite vectorial competence.

Other modes of transmission have been reported, such as vertical and sexual transmission in humans, and the potential role of other mosquitoes or natural reservoirs have been raised. Multiple Zika cases in different countries were reported to be individual cases of sexual transmission of Zika virus [32, 33, 34, 35]. We observed a much higher number of Zika cases in women (17,585 cases) than in men (7,628 cases) in the city of Rio de Janeiro. Coelho *et al.* [36] describe higher incidence of Zika in adult women in the city of Rio de Janeiro, but multiple causes could account for a disproportionate number of cases among women, such as awareness due to pregnancy-related risks or even a bias in symptomatic cases. Yakob *et al.* [37] argue for a low risk of sexual transmission based on model proposed for HIV transmission [38].

Given our current knowledge regarding the entomological parameters of ZIKV transmission, potential *R*_0_ for Zika assessed by only such parameters and dengue’s reproduction number should be about 1.4 times greater than the reproduction number of dengue. This result is in agreement with the estimates obtained from notification data. Therefore, we believe this approach can be used to evaluate the potential risk of Zika in areas with recurring epidemics of dengue. This method can be used as a risk assessment tool by Public Health authorities to inform disease mitigation policy.

## Acknowledgements

We would like to thank Dr Claudio Struchiner and Dr. Marília S. Carvalho for their helpful comments during the development of this study. We also acknowledge CNPq and FAPERJ for financial support.

## Author contributions statement

L.B., O.C., D.V., L.C., M.G., F.C. and C.C. conceived the analysis, O.C., B.D., M.L. and V.S. gathered the data, L.B, D.V. and L.M. developed the computational code. All authors analysed the results and reviewed the manuscript.

## Additional information

**Competing financial interests** The authors declare no competing financial interests.

